# Utilizing large language models to construct a dataset of Württemberg’s 19th-century fauna from historical records

**DOI:** 10.1101/2025.10.14.681982

**Authors:** Maximilian Teich, Belen Escobari, Malte Rehbein

## Abstract

Constructing datasets on past biodiversity from historical sources is crucial for understanding long-term ecological changes. Typically, compiling such datasets relies on prior knowledge of the sources’ composition and requires considerable manual effort. To overcome these challenges, we implement an automated approach based on prompted large language models (LLMs) to detect mentions of species in texts from 19th-century Württemberg and link these mentions to identifiers in the GBIF database. Based on our evaluation, we find that LLMs can reliably identify species in the texts with high recall (92.6%) and precision (95.3%), while providing estimates of the correct species identifier with considerable accuracy (83.0%).

## Introduction

Ecosystems and the changes they undergo can only be comprehensively understood and assessed in light of their past. Estimates of human influence on ecosystems, explanations for the extinction of certain species, or the formulation of realistic goals for conservation efforts all rely on knowledge of past developments. Historical ecology [1, 2, 3, 4, 5, 6] has established itself as a “framework for studying studying the past and future of the human–environment relationship”[7]. A central problem of historical ecology—as in all historical study—is the impossibility of directly observing the past. Instead of such observation, one must rely on the often incomplete historical record to draw conclusions about past circumstances. This is especially challenging as many available data sources are scarce or cover only small time frames or geographic areas. As Vellend et al. [9] note, this often necessitates the employment of unconventional methods and a creative use of data sources which may not originally have been intended to store information on past ecosystems.

Reconstructing datasets on past flora and fauna from fragmentary sources is a central part of historical ecology, ensuring that conserved information pertinent to present questions can actually be accessed and used by researchers, rather then being lost to history. The addition of new digital methods promises to help with this problem. Recently, Rehbein [10] has described these efforts as part of Computational Historical Ecology, in which “Natural Language Processing, Machine Learning, and Geospatial Analysis would support the extraction, classification, and integration of ecological information from a wide variety of archival materials.” The development and evaluation of approaches integrating such methods is ongoing.

Previous research has addressed the problem of constructing fauna datasets from textual historical sources in various ways: Some studies extract the relevant information largely by hand, as Turvey et al. [11] did for Gaboons from Chinese gazetteers, Clavero for fish species in 16th-century Spain, or Govaerts [13] for birds in 14th-century Holland from financial records. Working with structured data in form of tables, Rehbein et al. [14] designed a digitization workflow to build a dataset of 44 species documented in an 1845 survey of Bavarian wildlife. Blanco-Garrido et al. [15] mined geographical dictionaries for species names to derive a dataset of freshwater fish in 19th-century Spain, as did Viana et al. [16] for multiple Iberian species. Clavero et al. [17] combined text queries with spatial modeling to estimate historical Spanish wolf populations. All these approaches relied either on highly structured source data or substantial manual effort in compiling the final datasets or in providing training data for more automated solutions.

In this study, the fauna of the southern German kingdom of Württemberg during the 19th century is examined. Information on animal sightings from the time was collected in a series of government-issued descriptions of the kingdom’s districts. Presented in prose form, these texts contain accounts of the fauna, written without a uniform structure. To generate a dataset of all animals described in the source material and to assess its quality, solutions based on large language models (LLMs) were implemented and evaluated for the following tasks: First, all mentions of animals in the texts were recognized, regardless of textual structure, language (usually Latin or German), or spelling variations, a task generally known as Named Entity Recognition (NER). Second, these tokens were used to unambiguously identify the documented animal species by linking the tokens to a scientific authority file.

We demonstrate that datasets on past fauna can be constructed in a highly automated fashion from unstructured sources using LLMs, without fine-tuning and thus without the need for dedicated training data. We show that species detection from historical texts can be achieved with high recall and precision, despite challenges posed by historical language. By comparison, linking detected species to authority files is more error-prone but still provides useful indications of a species and its taxonomic classification.

In conclusion, we consider this approach a valuable option for building datasets from historical text sources, particularly when the scope of the material prohibits manual processing.

## Materials and methods

### Source Material

This study utilizes texts taken from 19th-century regional studies on the 64 *Oberämter* (administrative districts) of the southern German kingdom of Württemberg (see Figure 1). This kingdom had been formed only in 1806 under French hegemony and maintained its status after the political re-organization of 1815. Territorial changes during the Napoleonic era and the unavailability of reliable land registers motivated the government to begin a systematic land survey of the kingdom in 1818 [18]. By 1820, a Royal Statistical Office was established to support this survey. In particular, the office was instructed to record the geography, history, culture, and natural history of the various districts. This establishment was originally driven by the hopes that a better understanding of the newly integrated districts might help the royal administration increase tax revenues; thus the office was placed under the auspices of the minister of finance [19]. However, this purpose was soon eclipsed by that of national integration. Since the population of the new kingdom had a variety of cultural and political traditions that were not necessarily compatible with the idea of a unified monarchy, the office and its publications soon were instrumentalized to construct a common Württembergian national identity and to promote Württembergian patriotism [19].

**Fig 1.**
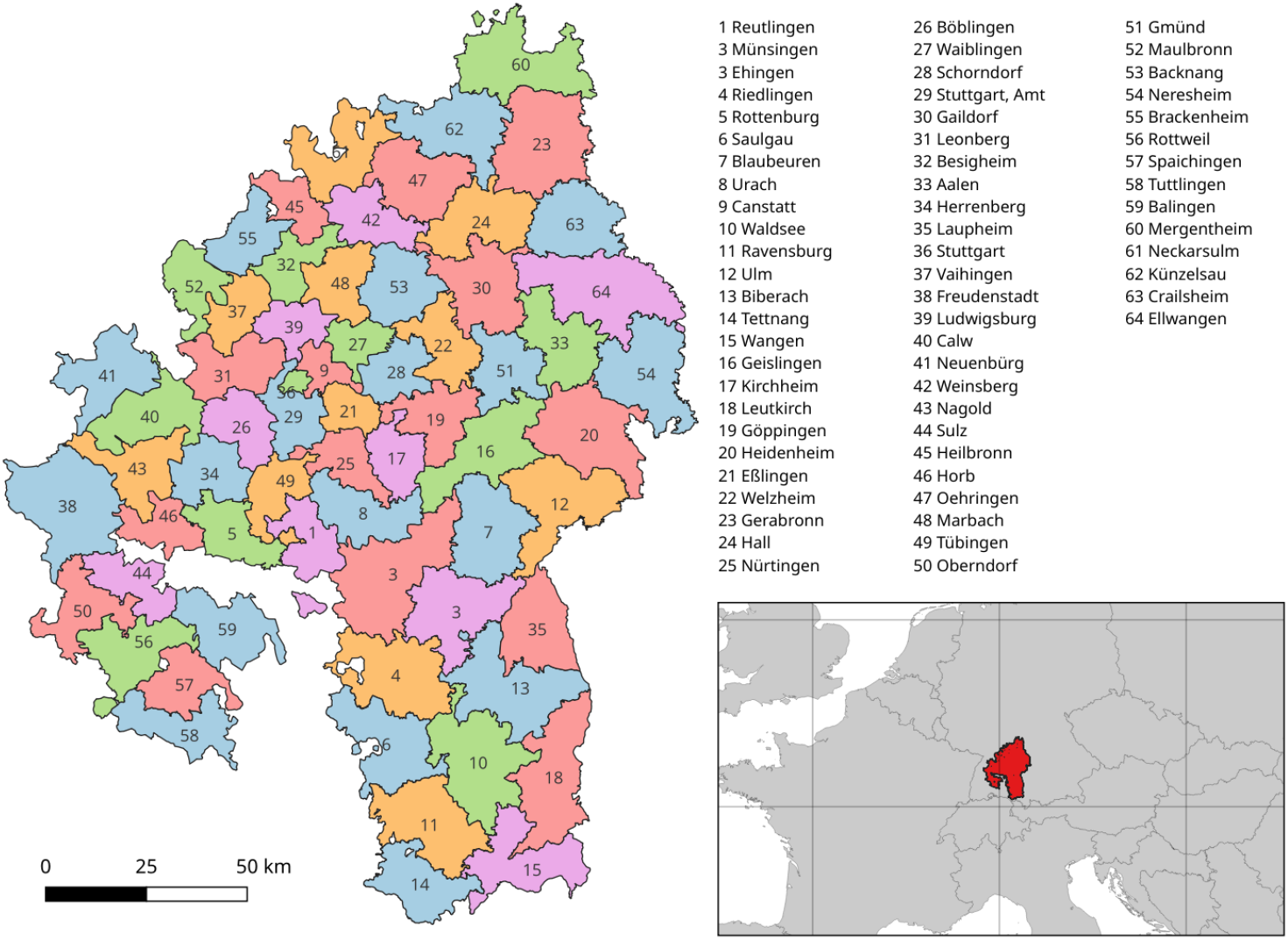
Districts. The districts of the kingdom of Württemberg in 1848 and the position of the kingdom in Europe with present-day borders shown for reference (low right).

The geographer and statistician Johann Memminger, the office’s first leading figure [20], advocated regional studies as a means for national integration [21]. In the preface to his 1822 yearbook on Württembergian statistics, he stressed that “there is no Württembergian people yet; every part is a stranger to the others”. He hoped for a “vitalization of national spirit” through knowledge of the fatherland [22]. Hence, the office’s work should not be seen as merely academic. In light of its political motivation, its projects also appear as attempts to educate the kingdom’s population on the surrounding regions and to instill a sense of unity.

Between 1824 and 1886, the Royal Statistical Office published 64 volumes of so-called district descriptions, each dedicated to the description of one of the kingdom’s districts and typically released at a pace of about one per year. Over several decades, these volumes grew more ambitious in scope; whereas the first volume had 158 pages, the last one was published in two parts with a total of 883 pages. Initially, Memminger compiled the first volumes based on his own research and information gathered from local contacts. The last volume in the series recounts these early efforts by Memminger and his assistant and the gradual expansion of the project [23]. As their production became increasingly professionalized, the editors could rely on a network of civil servants and part-time contributors with higher education [24]. Unfortunately, the texts generally do not include citations that identify the specific sources used by the editors. However, the original materials used to compile the texts are still preserved in the archives of the statistical office of Baden-Württemberg and could serve as a basis for future research into the volumes creation [25]. At least some of these records had been digitized by 2022 (description of the collection available online from the statistical office.) The volumes themselves are available as scans through various public libraries and Google Books. A Wikisource project has gathered these publicly available sources and manually transcribed all volumes into machine-readable texts (Wikisource.) These data are used in this study.

A notable feature of the district descriptions is the inclusion of a chapter on the fauna of each respective district, providing valuable historical insights into regional biodiversity. These chapters serve as the foundation for this study, offering a wealth of textual information. However, the quality of the fauna chapters varies greatly between the different volumes. While some merely note the absence of special or remarkable animals (e.g. in Kirchheim 1842, Künzelsau 1883, Crailsheim 1884), others provide detailed accounts listing hundreds of species. In some cases, the authors also discuss the significance of specific animals for local economies or cultural traditions. For example, the volume on Neckarsulm (1881) relates details on species of fish caught in the district, their culinary appeal to the local population, and special techniques for catching barbels (*Barbus fluviatilis*) with tridents in wintertime. The lengths of the chapters range from a brief 18 words for the rural Künzelsau district to a comprehensive 9682 words in the well-studied university town of Tübingen (1867), with a mean chapter length of 885 tokens and a general trend toward longer texts in later publication years. In total, by extracting the fauna chapters from each volume, a data set of 56,640 tokens was created.

### Entity Recognition

The first task is the recognition of n-grams naming animal taxa as presented in the chapters. This task also includes collecting information on the presence of a given animal, as some species are often recorded by the authors as being absent or extinct.

Previous research has looked into similar entity recognition problems in various ways. In cases where the texts have a predictable structure, such as in tables, encyclopedic entries, or uniform paragraphs, entities can often be identified by searching for specific patterns or strings [26, 27, 28, 29]. For texts without such structure, some authors have applied machine learning [30, 31, 32]. Several datasets have been published to help evaluate such approaches, but are not tailored to specific historical settings [33, 34, 35, 36, 37, 38]. In addition to traditional machine learning, fine-tuned language models have been integrated into workflows for species recognition [39, 40, 41]. Ehrmann et al.[42] offer an overview of all these approaches and outlines the challenges commonly encountered when dealing with named entity recognition (NER) in historical documents. More recently, research has focused on the use of prompt-based LLMs for retrieving named entities from historical texts without additional training data. While Tudor et al.[43] highlights the difficulties posed by hallucinations and Gonzales [44] finds this approach less effective than fine-tuned neural models, Hiltmann et al.[45] shows that it can substantially outperform traditional NER frameworks that lack training data, and emphasizes its potential benefits for historical research. In the realm of biology, Gougherty and Clipp [46] demonstrate how LLMs can be used to reliably extract information on the spread of pathogens from the literature, as does Scheepens et al.[47] for invertebrate pests and pest control agents. In a previous study, we looked at identifying common problems in the detection of species names in historical texts by using a small sample of the district descriptions, without attempting to process the entire corpus ([48]).

The following challenges to solving the entity recognition task were observed when assessing the historical texts:

- First, the texts share no common structure. Some volumes tend to enumerate species names in long sentences, whereas others devote multiple sentences to details on a species. Hence, one has to account for all possible ways in which animals may be referenced. This can include names for individual species or names for an entire class of animals.
- Second, animals can appear under their German vernacular names, scientific names, or both, which is not consistent across volumes. For example, the fire salamander (*Salamandra maculosa*) is noted in Backnang (1832) as “der gefleckte Erdmolch (Salamandra maculosa),” in Stuttgart (1851) just as “Salamandra maculosa,” in Oberndorf (1868) as “der Erdmolch” and in Spaichingen (1876) as “der gefleckte Salamander (Salamandra maculosa).” Therefore, any recognition method should be capable of determining when two different names refer to the same animal.
- Third, the names themselves often deviate from modern spellings. For example, the kestrel *Falco tinnunculus* is usually given as “Thurmfalke” instead of the modern “Turmfalke”, the burying beetle *Nicrophorus vespillo* as “Todtengräber” instead of “Totengräber.” Some historical vernacular and scientific names are obscure or no longer in common use. Scientific names are also frequently abbreviated. These problems may render them unintelligible even to domain experts.

Considering these multiple challenges and the general absence of task specific training data, a zero-shot approach using large language models (LLMs) could be a viable solution.

To allow LLM-based recognition, the historical texts were first split into chunks of 200 tokens, which proved to be a manageable size in the subsequent process. An overlap of 50 tokens between the chunks ensures that each animal name is included in full in at least one chunk. This process resulted in a total of 934 chunks. Each chunk was then submitted to an LLM following a prompt specifying the task: to list all animals mentioned in the historical text by both their vernacular and scientific names, if provided, and to indicate whether each animal is described as present or absent. The prompt also included instructions to return the results as a JSON element including the vernacular name, scientific name, and presence information, for example about the European goldfinch (*Carduelis carduelis*):

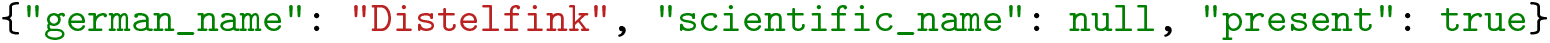

This format requirement was enforced through a validation loop, which repeated the prompt until a correctly structured response was received.

As a means of evaluating the quality of responses generated by different LLM and prompt variations, a test dataset was created. For 50 randomly sampled text chunks, we annotated all names and information about the animals’ presence using the same JSON format. We used the same spelling as the original historical texts and assumed an animal to be present when no absence was implied. This annotation yielded data for 435 mentions of animals in the sample. Using these test data as ground truth, the LLM responses were evaluated using three metrics:

1. A recall score showing how many of the animals found by the human annotator were also recognized by the LLM.
2. A precision score, indicating how many of the animals found by the LLM were correct, i. e. also annotated by the human.
3. An additional precision score for the information on presence, showing in how many cases the LLM agreed with the human annotator on whether an animal is described as being present or absent.

This proved challenging, as all tested LLMs tended to modernize historical spellings in their outputs. Additionally, the texts often used abbreviated scientific names, which the LLMs attempted to fully spell out. To address these issues, we applied two different matching strategies to compute the metrics, following the approach of Gonzales-Gallardo et al.[44]:

1. A strict matching function, which only considers items a match if the vernacular or scientific names are exact string matches.
2. A fuzzy matching function, which allows for minor variations (an editing distance of 2 was set as acceptable) and assumes that a match is still valid if the first part of a scientific name is abbreviated in either the human’s or the LLM’s output.

Testing three common LLMs and adjusting the prompt for optimal performance, we found that the models can generally detect the majority of documented species in the texts. GPT-4o performed best on the task, followed by the Gemma 2 (see Table 1). As one would expect, the metrics’ values are higher when fuzzy matches are considered, giving the best model a recall score of 93% and a precision score of 95%. Under strict matching conditions, the recall and precision scores drop to 83% and 84%, respectively. Regarding the presence or absence of animals, all models score very high. As the calculation of this metric can only consider the matches found between the Ground Truth and the solutions of the LLMs’, there is no discernible difference between the two matching functions. Generally, the best performance of an LLM is comparable to that reported for tools tuned on training data to identify taxa [36, 38].

**Table 1.**
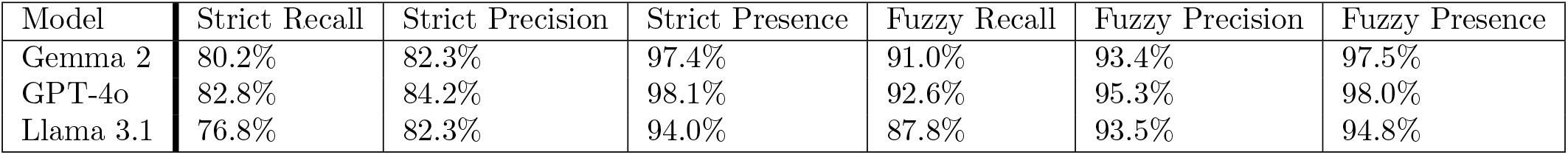
Evaluation metrics for tested LLMs on entity recognition task. Edit distance for fuzzy matching is set at 2.

| Model | Strict Recall | Strict Precision | Strict Presence | Fuzzy Recall | Fuzzy Precision | Fuzzy Presence |
| --- | --- | --- | --- | --- | --- | --- |
| Gemma 2 | 80.2% | 82.3% | 97.4% | 91.0% | 93.4% | 97.5% |
| GPT-4o | 82.8% | 84.2% | 98.1% | 92.6% | 95.3% | 98.0% |
| Llama 3.1 | 76.8% | 82.3% | 94.0% | 87.8% | 93.5% | 94.8% |

### Data Linking

Building on the extraction of vernacular and scientific names from the texts, a second task involves linking these names to records in an authority file. By mapping the names to such unique records, historical mentions of animals can be disambiguated, and additional information on each taxon can easily be added for further analysis. The *Global Biodiversity Information Facility* (GBIF), an international network and data infrastructure project which aggregates data on taxa from a wide range of sources, was chosen as the target authority file. GBIF assigns a unique identifier to each taxon and records all names associated with it along with other information such as distribution data. For each mention of an animal in the historical texts—whether it is given by a scientific name, a German vernacular name, or both—the corresponding identifier in GBIF must be found.

Two approaches were considered to complete the task. The first involved matching tokens identified in the historical text against the GBIF database using its API (GBIF API Reference.) This API allows for the search of scientific names, supports fuzzy searches, and returns confidence scores for uncertain matches. The separate API function for searching vernacular names is problematic, as this is implemented as a simple text search. This function can return irrelevant results from the database’s resources. For example, a search for the German vernacular name “Bär” (*Ursus arctos*) may also return other species described in the literature by the botanist with the same surname: Johannes Bär. To avoid this ambiguity, the API was bypassed by downloading parts of the GBIF database and performing local queries. As a basis, a dataset containing approximately 12,500 German vernacular names (compiled by Holstein and Monje [49]) was used. This dataset includes names of animals and plants from southern Germany and is itself integrated into GBIF. The data were further enriched by retrieving all synonymous vernacular names listed in GBIF for each record. Historical vernacular names were then matched against this dataset to identify the corresponding GBIF entries. However, the direct lookup of names is still prone to several issues: historical names often exhibit spellings diverging from modern conventions, and scientific names are frequently abbreviated. Additionally, some historical names may have fallen out of use or are too rare to have found entry into GBIF. In such cases, one cannot expect to find corresponding matches in the dataset. Hence, the results of this approach will generally have a low recall at a high precision.

The second approach to matching historical names and authority records employs LLMs. Previous research has explored the use of prompt-instructed LLMs for entity matching tasks. Farrell et al.[50] suggest using such models to harmonize data directly from textual sources. Peeters et al.[51] find them to be effective in applications where no training data are available or where many unseen cases are expected, as is the case in our task. However, quality of the results depends on the extent of knowledge represented within the models. Dorm et al.[52] and Elliott and Fortes[53] tested the knowledge of popular LLMs regarding geographical species distribution, showing varying results which likewise suggest a strong dependence on the models’ training data. Castro et al. [54] used LLMs to identify species in research papers and newspaper articles with a few-shot approach and also found that the models tested vary widely in their ability to perform the task. As we must assume that common LLMs are not necessarily trained on 19th-century German biological texts, we devised prompts in a way to pass both the historical names and the original text chunk to an LLM, to ensure that there is some context available. Based on these inputs, the model is instructed to suggest a modern scientific name for the animal in question. Initial experimentation showed that this LLM-based approach yielded less precise results than the pure database lookup approach, even with the contextual information offered by the text chunks. At the same time, it achieved perfect recall, as the LLM would always return at least a plausible result.

Given the shortcomings of both approaches, we designed a combined workflow for optimized performance on the task: In a first step, the extracted scientific and vernacular names are looked up in the locally stored copy of the database. Only cases where no match is found are passed to a second step, in which an LLM is prompted to provide the animal’s modern scientific name based on the historical names and contextual information. This result can then be used to attempt another lookup in the GBIF database (see Figure 2).

**Fig 2.**
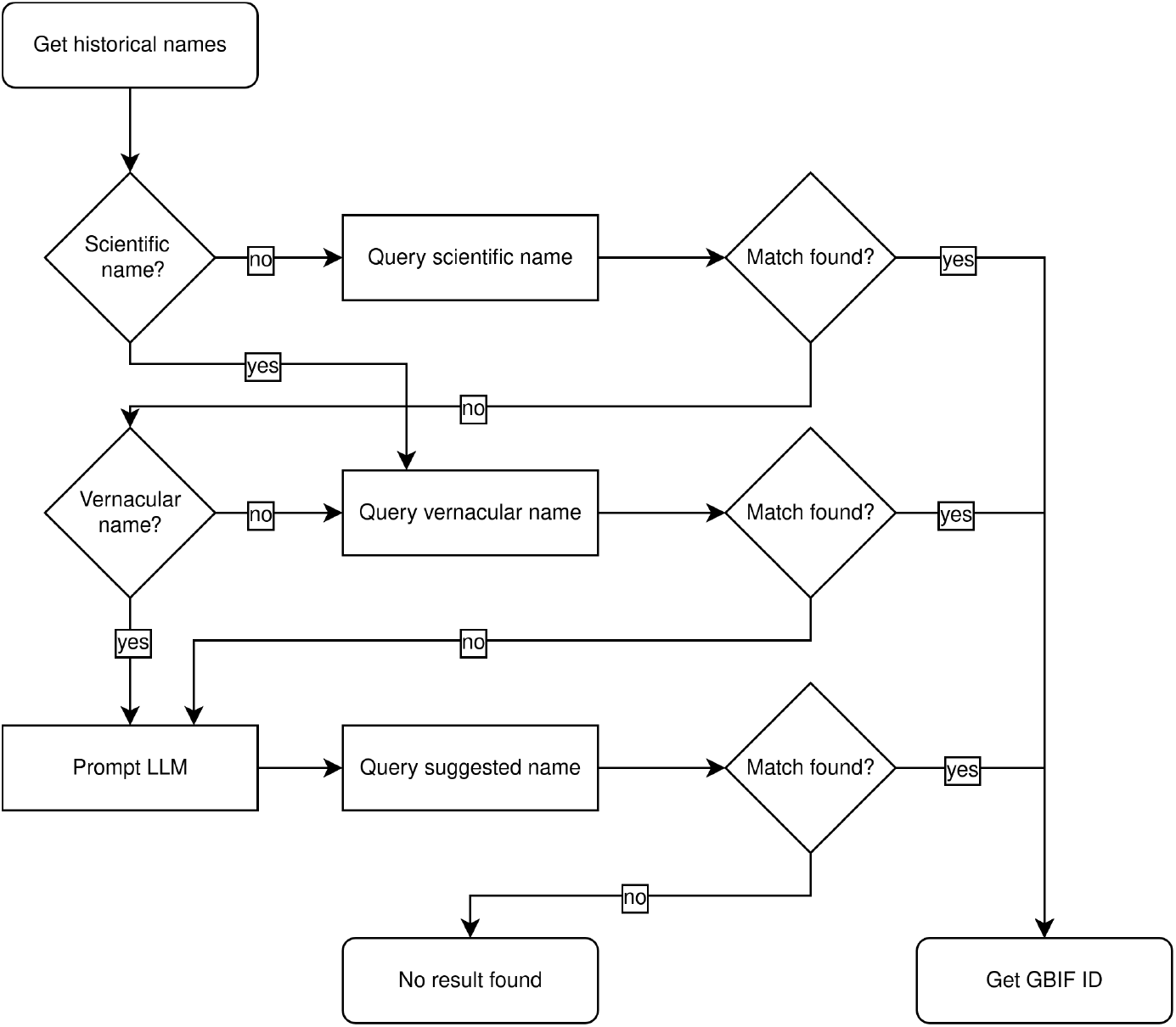
Worflow. Combined approach workflow using direct lookup of names in the GBIF database and an LLM in case historical names are not found directly.

To evaluate the quality of the results produced by this combined approach, an additional test dataset was manually created as a ground truth. For 300 randomly selected triples consisting of scientific names, German vernacular names, and the original text chunk as context, initial experimental results were evaluated by a domain expert. The expert also recorded the correct solution for each case. However, in 27 of the 300 test cases (9%), even the expert could not determine the correct solution with certainty. For example, for Freudenstadt (1858) authors documented “Schaumwurm (Cercopis spumaria).” This scientific name is a synonym of *Philaenus spumarius* (common meadow spittlebug), but neither the common name, nor the historical synonym is recorded in GBIF or any other information source. These uncertain cases were excluded from the accuracy calculation but suggest a noteworthy baseline error for the task. After the evaluation, all synonyms of the correct solutions available were queried from GBIF to collect all possible valid answers. Against these lists of valid answers, both the GBIF lookup results and the LLM outputs could be automatically evaluated. An accuracy score was introduced to indicate how often the combined approach matched the solution of the domain expert. A recall metric indicates for how many cases a solution was found.

Results of the evaluation are shown in Table 2. Using only the direct lookup of historical names in the GBIF database, the results of the domain expert could be reproduced in 92% of the test cases. However, this approach alone only yields results in 61% of cases and returns no result in the remaining 39. When an LLM is included to find solutions for these remaining cases by suggesting modern equivalents to the historical names not found directly in the database, all of them were solved. However, with the introduction of the LLMs, the overall accuracy drops, as the suggested names are less likely to conform with the expert’s solution. The best performance was achieved by using Llama 3.1, with an accuracy of 83% with the remaining inaccurate results often close to the true solution. For example, the “Saatgans” (*Anser fabalis*) is falsely linked to the “Graugans” *Anser anser*, or the “Birkhuhn” (*Tetrao tetrix*) is mistakenly for the British subspecies *Tetrao britannicus*.

**Table 2.** Evaluation metrics for the combined approach to the data linking task.

| Model | Recall | Accuracy |
| --- | --- | --- |
| Gemma 2 | 100.0% | 77.0% |
| GPT-4o | 100.0% | 80.7% |
| Llama 3.1 | 100.0% | 83.0% |
| without LLM | 60.9% | 92.0% |

It should be noted that the performance of the task depends on the kind of test case observed. This can be seen when grouping the cases in the test dataset by the taxonomic class and evaluating each class separately (see Table 3). The largest group in the dataset are birds, accounting for 139 of the test cases. For this group, Llama 3.1 achieves an accuracy of 87%. Amphibians (n=11) and Mammals (n=50) show the best results with accuracy scores of 91% and 88%, respectively, also with Llama 3.1. At the other end of the spectrum, insects (n=23) can only be linked correctly in 48% of cases.

**Table 3.**
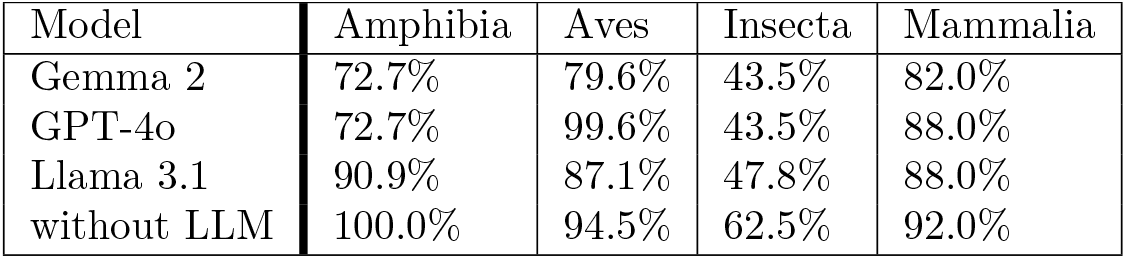
Accuracy results by species class.

| Model | Amphibia | Aves | Insecta | Mammalia |
| --- | --- | --- | --- | --- |
| Gemma 2 | 72.7% | 79.6% | 43.5% | 82.0% |
| GPT-4o | 72.7% | 99.6% | 43.5% | 88.0% |
| Llama 3.1 | 90.9% | 87.1% | 47.8% | 88.0% |
| without LLM | 100.0% | 94.5% | 62.5% | 92.0% |

Considering these variations in the performance of the workflow, the evaluation was further extended to additional taxonomic levels: For each element in the test set, GBIF entries on phylum, class, order, family, and genus were queried. Non-empty values were then compared to those for the outputs of the various LLMs to evaluate performance on each level. The results are shown in Table 4. For phylum and class, all LLMs produced nearly perfect results, with errors no greater than about 2%, even at the order level. However, performance noticeably declined at the family level, dropping to an accuracy of only 89% for the best-performing Llama 3.1 model at the genus level. These findings suggest that most of the method’s errors arise from confusion at the lower taxonomic levels—family and genus. Nevertheless, the resulting information about an animal’s order, class, and phylum is likely to be accurate, even if the method fails to link to the correct species identifier.

**Table 4.** Accuracy results for other levels of the taxonomy.

| Model | phylum | class | order | family | genus |
| --- | --- | --- | --- | --- | --- |
| Gemma 2 | 99.2% | 99.2% | 98.1% | 93.2% | 87.2% |
| GPT-4o | 99.6% | 99.6% | 97.7% | 94.0% | 88.2% |
| Llama 3.1 | 99.6% | 99.6% | 98.1% | 92.5% | 88.9% |
| without LLM | 99.4% | 99.3% | 99.4% | 98.1% | 96.3% |

Lastly, we evaluated the performance of the LLMs with respect to the diachronic nature of sources. Considering the 62 years time span in which the documents were written, we hypothesized that animal names became more standardized over time and approached modern conventions, which should lead to better model performance for more recent texts. To account for the uneven distribution of the data over the years, the test dataset was divided evenly into four quarters of equal token size. The LLMs’ outputs were then evaluated separately for each quarter. Contrary to our expectation, the results shown in Table 5 do not indicate a trend toward more easily interpretable names over time. While the best-performing Llama 3.1 model scores highest on the most recent quarter of names from the 1870s and 1880s, it shows its worst performance on the second most recent quarter. The other two models achieve their best results on the second quartile, which contains names from the 1850s and 1860s. Overall, the diachronic analysis of the LLMs’ performance yields no significant results.

**Table 5.** Accuracy results for four time spans.

| Model | 1825-1852 | 1853-1863 | 1865-1871 | 1872-1886 |
| --- | --- | --- | --- | --- |
| Gemma 2 | 72.6% | 80.1% | 78.3% | 76.2% |
| GPT-4o | 81.4% | 83.8% | 78.2% | 79.3% |
| Llama 3.1 | 82.9% | 82.3% | 79.7% | 87.3% |

## Results

A dataset of the animals documented in the district descriptions was built using the entity recognition workflow with GPT-4o and the data linking method with Llama 3.1 (see S1 Dataset). Additionally, the dataset includes information created by employing only a database lookup, without using an LLM for linkage. This is intended for applications that require higher precision, at the expense of significantly lower recall of the animals mentioned in the original texts. In both cases, duplicates where animals are mentioned multiple times within a district were removed, as these typically result from overlapping text chunks.

The dataset, which is expected to capture the vast majority of animal mentions from the original texts, includes 6794 entries. Figure 3 shows how these entries are distributed between districts and publication years. In three districts, no animal names could be detected. These results are correct in so far as the corresponding texts state that no research was conducted on the fauna of the district (Kirchheim) or that the authors stated that the fauna does not differ noticeably from that of a previously described neighboring district (Crailsheim and Künzelsau), thus referring to the other volumes. For the remaining districts, the number of detected animal mentions ranges from two in Cannstatt to 499 in Tübingen, with a mean of 106 mentioned animals. In general, the number of mentions increases over time. Simultaneously, the overall length of the fauna-sections normalized to the number of animals documented decreases in the later volumes, suggesting that the texts became less descriptive over time and began to favor enumerations.

**Fig 3.**
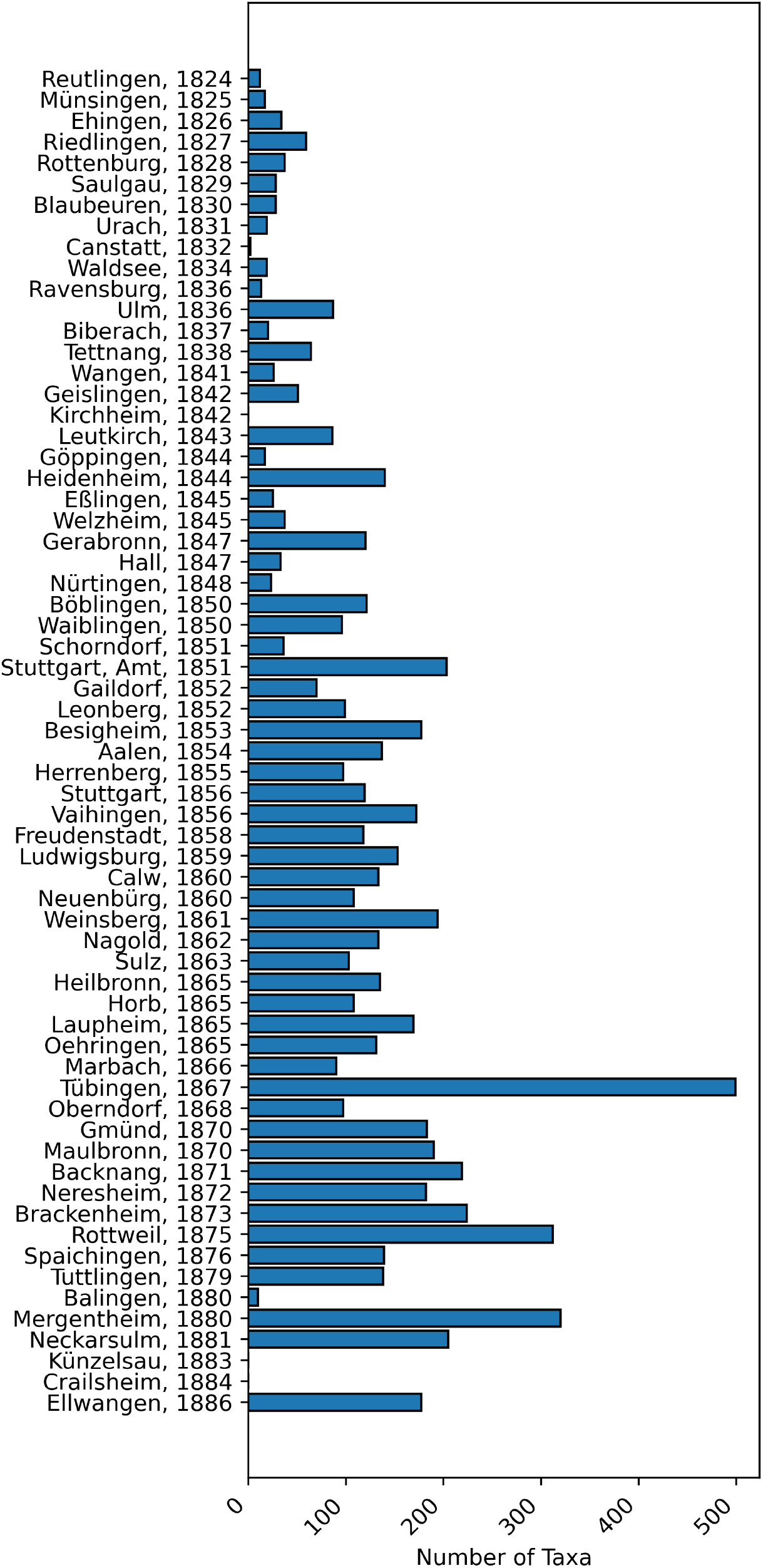
Taxa distribution. Distribution of detected taxa over districts and years.

In addition, the distribution of animal classes over time can be assessed within the dataset. Birds are mentioned most frequently overall, comprising 45.3% of the records, followed by insects (15.9%) and mammals (10.8%). For 11.5% of the animals found, there is no class entry in GBIF; based on the test dataset, these cases can be assumed to be almost exclusively fish, which are generally not assigned a class value. Figure 4 illustrates how the class values are distributed across the corpus. The share of birds remains high throughout and increases in the later texts. Both, the proportions of mammals and animals without a class value decrease over time, whereas the proportion of insects grows. The relatively smaller shares of less common classes—such as amphibians, gastropods, and squamates—remain relatively stable.

**Fig 4.**
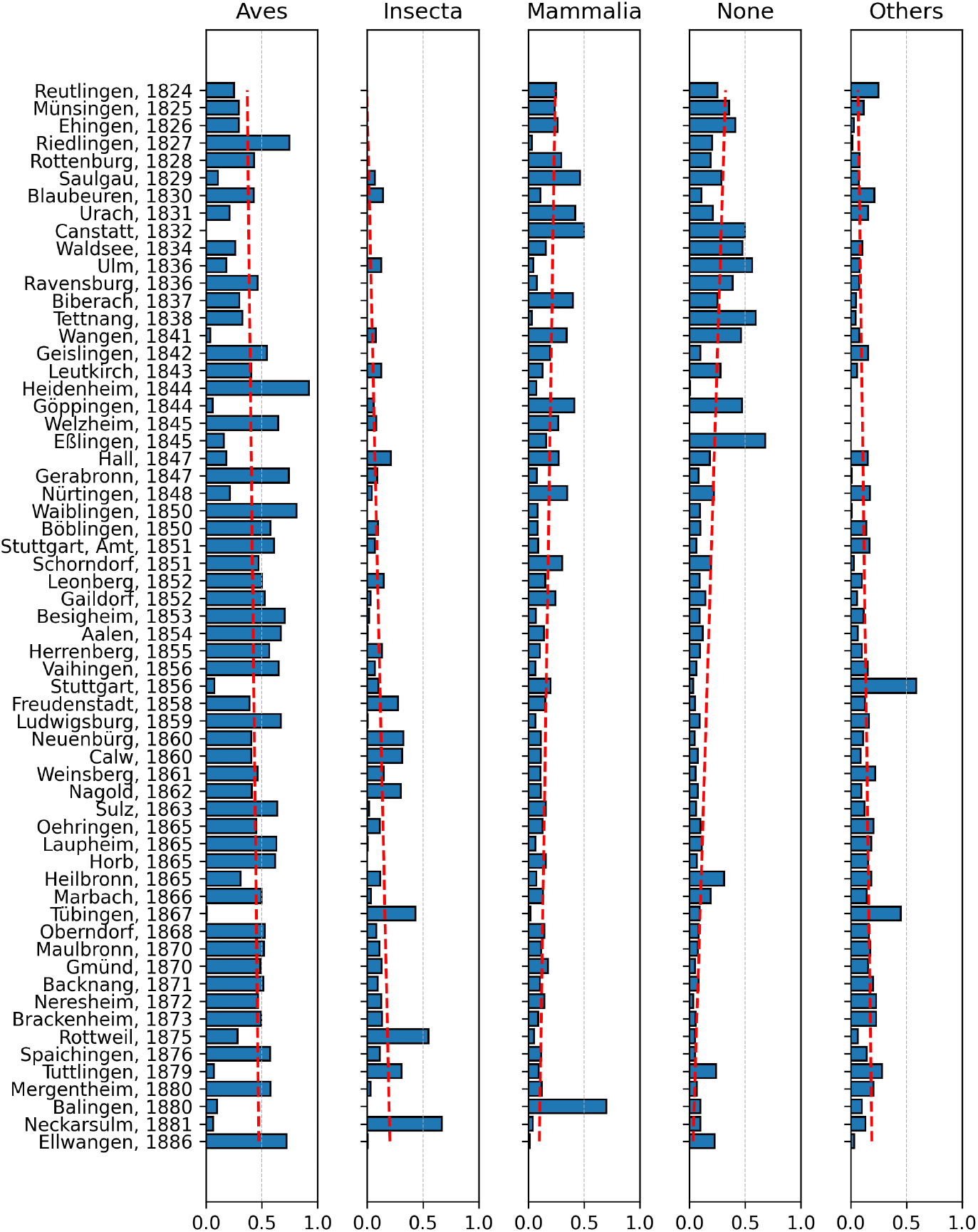
Class proportions. Proportions of most frequent classes of animals by districts and years. Regression lines in red indicate trends.

To gain a better understanding of the spatial distribution of the data, it can be projected onto a map of the districts. Since there is no readily available vectorized geodata, we created a custom GeoJSON file of the borders of the districts. This is based on a publicly available mid-19th-century topographical map of Württemberg at a scale of 1:200,000, produced by the Royal Statistical Office ([55]). The map shows the district borders in 1848, during the time the district descriptions were edited and published. As the borders were rarely and only slightly adjusted after 1810 [56], we assume the map gives a good representation of the districts extent. This historical map was georeferenced and retraced using the open-source QGIS software (see S2 GeoJSON).

Figure 5 visualizes the number of animals mentioned per district on the resulting map. It clearly shows that the central and southern districts—surveyed earliest—mention fewer animals. Conversely, the volumes for more remote districts in the west and north, generally published after 1850, contain more mentions. The university town of Tübingen (published 1867) stands out clearly as an outlier, suggesting it had been studied more intensively than any other district in the kingdom.

**Fig 5.**
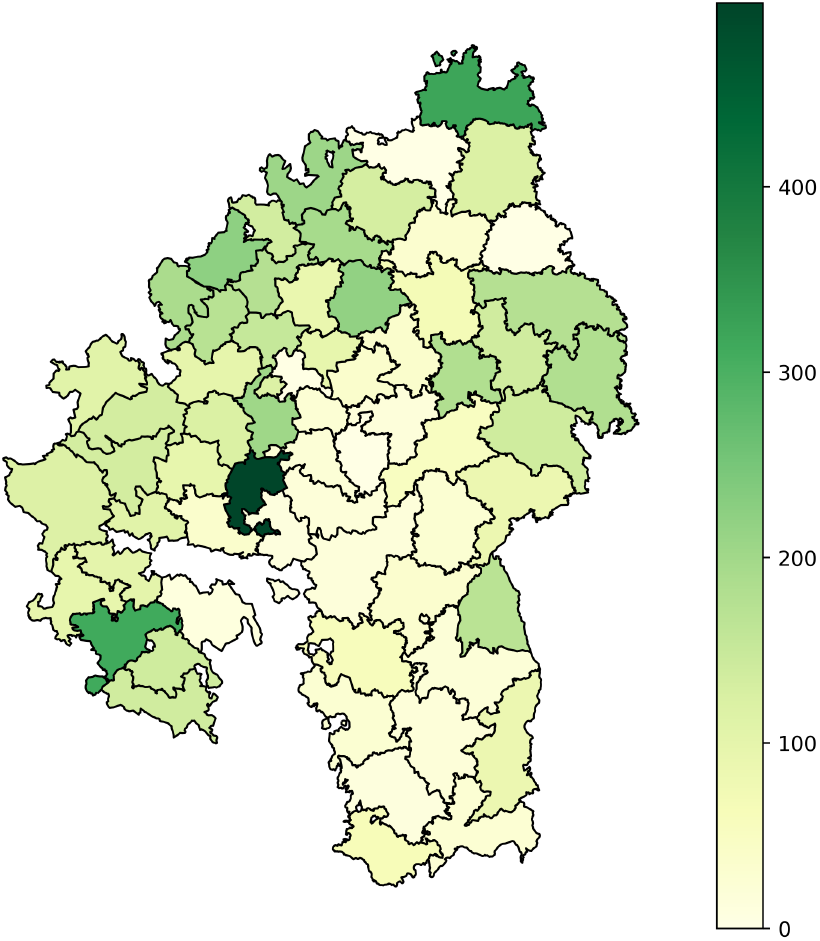
Animals by district. Number of extracted animal mentions by district.

The same visualization approach can be used to identify where individual species were documented. Even shorter texts usually document mammals that were common game animals. The latter are often described in the greatest detail and tend to be mentioned even when extinct in a district, sometimes with information on when they were last spotted or shot. For example, Figure 6 shows mentions of wild boar (*Sus scrofa*), roe deer (*Capreolus capreolus*), and hare (*Lepus europaeus*). The wild boar is frequently documented in early volumes, albeit only in the context of its extinction in the course of the 19th century. The later volumes largely omit references to wild boars, perhaps because their absence had come to be expected. Only four cases indicate the presence of wild boars and the respective texts explicitly describe them as rare and migrating from beyond the kingdom’s borders. In Mergentheim, the text even recounts in detail two boars being shot in 1870 and another spotted by a hunting party of military officers, highlighting how remarkable these occurrences were. In contrast, roe deer and hares are generally assumed to be present. Deviations from this expectation are often explained in the texts, such as for Maulbronn (1870), where the authors note that “the number of hunting enthusiasts is increasing by the day” and this had—along with illness—decimated the local hare population.

**Fig 6.**
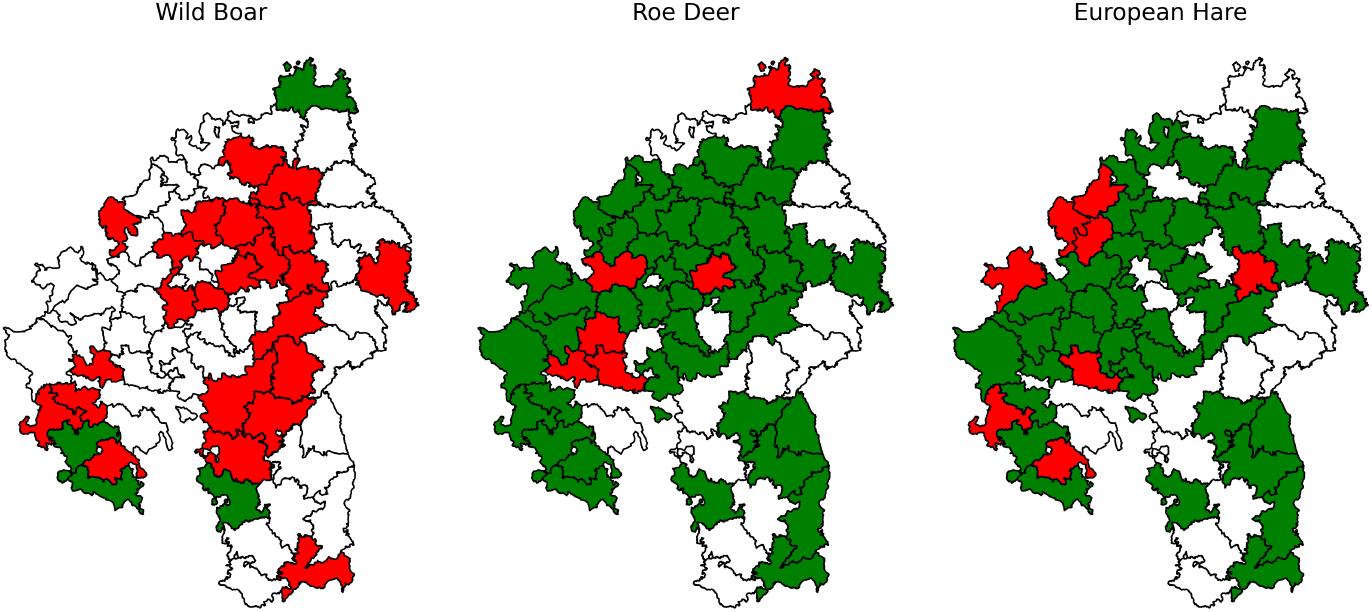
Animal Presence. Presence of wild boar, roe deer, and European hare in the districts. Green colour indicates presence, red absence, white no data for the district.

## Discussion

The problem of generating a fauna dataset for the Kingdom of Württemberg from unstructured 19th-century source texts in an automated manner was addressed by breaking it down into two tasks: The first task involved detecting animals mentioned in the texts as named entities. This was challenging due to the lack of structure in the texts, the archaic language, and the general uncertainty regarding the expected taxa. It was found that processing the texts in chunks with a prompted LLM and enforcing a structured response led to reasonably good results when evaluated using strict string matching metrics (recall = 82.8%, precision = 84.2%). When more lenient, but arguably more appropriate, fuzzy matching criteria were applied, this yielded better results (recall = 92.6%, precision = 95.3%). This suggests that, while not perfect, an LLM-based approach can detect biological taxa even in unstructured historical texts.

The second task of linking the retrieved taxa to entities in an authority file proved more difficult. Due to the use of archaic names in the source texts, finding a correct match was not possible in a substantial number of cases, even for human experts. The combined approach employed here—first looking up the retrieved names in a database of known taxa, and then prompting an LLM to suggest a modern equivalent if that failed—worked reasonably well. Overall, this approach can be expected to yield a species-level accuracy of 83.0% based on the test data. While this still implies a relatively high error rate, it should be noted that the accuracy scores are better at higher taxonomic levels. When identifying the class or order of an animal, the approach achieved very high accuracy scores of 99.6% and 98.1%, respectively. This shows that the approach, even though it may not reach human-level quality in linking historical taxa to modern records, can produce results of predictably high quality.

Given these results, a dataset of known quality has been generated. This demonstrates, that prompted LLMs can be valuable for building workflows to generate biodiversity datasets from historical text sources in a highly automated fashion. In particular, an LLM-based approach offers several advantages: it requires no predefined structure of the source texts, no prior knowledge of the animals described, and no need for special training data or fine-tuning. Additionally, it demands little manual effort.

Compared to datasets created by human experts from historical documents, such as [13, 17], these advantages are remarkable. Particularly in the analysis of historical source texts, where large amounts of data and a requirement for niche expertise may render traditional approaches unfeasible, LLMs can offer a viable alternative. However, these advantages may come at the expense of dataset quality especially at the lower taxonomic levels, as shown by our evaluation. When choosing to implement such an approach, it is important to consider the trade-offs involved. If available resources are limited relative to the scope of the historical sources, the use of LLMs is likely a promising option. The same holds true in cases where only a general assessment of the data is needed—for example, when an imperfect recall of mentioned species or a broad analysis of order-level taxonomic distribution is sufficient.

Conversely, when data of nearly perfect quality are required, human oversight remains indispensable. In any case, it is advisable to implement a method for evaluating the quality of the resulting dataset in order to be able to estimate the extent of potential errors.

## Supporting information

**S1 Dataset**. File containing the constructed dataset. Available from: https://doi.org/10.5281/zenodo.17277241

**S2 GeoJSON**. GeoJSON file of the *Oberämter* (district) borders of the Kindgom of Württemberg in 1848.Available from: https://doi.org/10.5281/zenodo.17277241

